# Telomere-to-telomere genome assembly of the clubroot pathogen *Plasmodiophora brassicae*

**DOI:** 10.1101/2024.04.03.587992

**Authors:** Muhammad Asim Javed, Soham Mukhopadhyay, Eric Normandeau, Anne-Sophie Brochu, Edel Pérez-López

**Author notes:** These authors contributed equally (Muhammad Asim Javed, Soham Mukhopadhyay).

## Abstract

Plasmodiophora brassicae (Woronin, 1877), a biotrophic, obligate parasite, is the causal agent of clubroot disease in brassicas. The clubroot pathogen has been reported in more than 80 countries worldwide, causing economic losses of hundreds of millions every year. Despite its widespread impact, very little is known about the molecular strategies it employs to induce the characteristic clubs in the roots of susceptible hosts during infection, nor about the mechanisms it uses to overcome genetic resistance. Here, we provide the first telomere-to-telomere complete genome of Plasmodiophora brassicae. We generated ∼ 27 Gb of Illumina, Oxford Nanopore, and PacBio HiFi data from resting spores of strain Pb3A and produced a 25.3 Mb assembly comprising 20 chromosomes, with an N50 of 1.37 Mb. The BUSCO score, the highest reported for any member of the group Rhizaria (Eukaryota: 88.2%), highlights the limitations within the Eukaryota database for members of this lineage. Using available transcriptomic data and protein evidence, we annotated the Pb3A genome, identifying 10,521 protein-coding gene models. This high-quality, complete genome of Plasmodiophora brassicae will serve as a crucial resource for the plant pathology community to advance the much-needed understanding of the evolution of the clubroot pathogen.

**SIGNIFICANCE:** Plasmodiophora brassicae (Woronin, 1877) is a devastating plant pathogen, member of the Rhizaria group, and is putting at risk the oilseed rape and cruciferous vegetable industry worldwide. Here, we present the first telomere-to-telomere genome of P. brassicae, and the first complete genome for a member of the Rhizaria taxon. We also provide a high-quality genome annotation featuring an enhanced gene model relative to previous studies. This complete genome will be of great interest to researchers focused on the ecology and evolution of the clubroot pathogen, as well as those studying other members of the Rhizaria group.

## INTRODUCTION

*Plasmodiophora brassicae* (Woronin, 1877) is a soil-borne protist that infects brassica crops, including the economically important canola (*Brassica napus*). By manipulating the host‘s hormone levels and cell division, the clubroot pathogen causes radical swelling of the root and hypocotyl, forming galls or club-like structures that affect the normal development of the host (Liu et al., 2020). As a biotrophic and obligatory parasite, the clubroot pathogen sequesters host nutrients to complete its life cycle, turning the roots into nutrient-sink galls that slows down its host’s growth compared to healthy plants (Malinowski et al., 2019). Since its discovery in commercial canola fields in Canada, the clubroot pathogen have cost hundreds of millions of dollars to the Canadian economy and the agriculture industry is still struggling to manage this devastating pathogen (Javed et al., 2023; Ochoa et al., 2023).

The clubroot pathogen belongs to the highly diverse, unicellular eukaryotic group Rhizaria, within the class Phytomyxea and the order Plasmodiophorids – a group of poorly studied plant pathogenic protists (Neuhauser et al., 2014; Mukhopadhyay et al., 2024). Due to the intracellular lifestyle and the soil-borne nature of Plasmodiophorid plant parasites *Spongospora subterranea, Polymyxa betae*, and P. *brassicae*, the extraction of high-quality DNA is challenging and often contains a high proportion of host DNA (Ciaghi et al., 2018; Decroës et al., 2019; Stjelja et al., 2019; Javed et al., 2023). Although efforts have been made to assemble a reference-quality P. brassicae genome, especially from several European groups, those are not complete, show discrepancies in their annotations and predicted number of gene models, and in some cases the data is not fully available to the scientific community (Schwelm et al., 2015; Stjelja et al., 2019; Daval et al., 2019; Bi et al., 2019; Li et al., 2023). Despite the identification and characterization of more than 40 clubroot pathotypes that pose a significant threat to the canola industry, high-quality, assembled, and annotated genomes for Canadian isolates remain unavailable, although some draft genomes exist (Rolfe et al. 2016; Sedaghatkish et al., 2019). Constructing a chromosome-level genome assembly of the clubroot pathogen is essential for advancing future research. This will enable more robust and cost-effective analyses in population and comparative genomics, facilitate the acquisition of functional insights, and help uncover structural variations among genomes of different clubroot pathogen isolates or pathotypes that can be linked to pathogenicity or evolution (Amezrou et al., 2024; Badet et al., 2024; Croll, 2024). Here, we present the first telomere-to-telomere genome assembly and *de novo* annotation of *P. brassicae*. To our best knowledge, this is also the first telomere-to-telomere genome assembly for any member of the group Rhizaria. This will serve as a pivotal reference genomic resource for the clubroot research community and to better understand protist biology.

## RESULTS AND DISCUSSION

### Telomere-To-Telomere Genome Assembly

High molecular weight (HMW) DNA was extracted from the resting spores of a field isolate, previously pathotyped as 3A (hereafter Pb3A). This isolate, which was initially collected in 2013 from canola fields in Alberta, Canada, was kept under controlled conditions in susceptible Westar canola ever since. A total of 20 μg of HMW DNA was used for PacBio-HiFi, Oxford Nanopore, and Illumina library preparation and sequencing.

Long-read sequencing using PacBio HiFi produced a total of 4 gigabases (Gb) of data, 6.22 Gb using Oxford Nanopore, while short-read sequencing using Illumina produced 16.9 Gb of data, with depths of coverage above 300x for all three technologies. We used both Pacbio and Oxford Nanopore long reads to produce an initial hybrid assembly which was further polished using the Illumina short reads. The assembly, produced with Hifiasm, generated 269 contigs. Out of those, the top 20 largest contigs were assigned as full-length chromosomes with intact telomeres at both ends, except for chromosomes 5, 9, and 10, where we couldn‘t capture telomeres at one end of each chromosome. To find the missing telomeres, we conducted further Hifiasm assemblies using longer PacBio HiFi (read length cut-off 3Kb) and Oxford Nanopore long reads (read length cut-off 8Kb). Additionally, we generated a RAVEN assembly using only Oxford Nanopore reads and subsequently polished with Illumina short reads. In the RAVEN-based assembly, chromosome 9 presented intact telomeres at both ends. The sequence of RAVEN generated chromosome 9 showed very high co-linearity with the Hifiasm produced chromosome 9. Consequently, we replaced chromosome 9 in our final assembly with this complete version. After aligning one of the Hifiasm hybrid assemblies (longer read length) with our final genome assembly using D-GENIES, we found two small contigs having sequence overlap with the ends of chromosome 5 and 10. These two small contigs contained the telomere sequences that were missing in the final alignment. Thus, these were added to the ends of their respective chromosomes. To validate whether this was an artifact, we mapped PacBio HiFi reads to all chromosomes. We observed continuous read coverage (>100x) spanning the regions where the fragments were attached. The telomere sequence TTTTAGGG (T4AG3) assembled in the Pb3A genome was previously reported in some chromosomes of P. brassicae strain e3 and other evolutionary-related organisms, including the green algae *Chlamydomonas reinhardtii* (Petracek et al., 1990) and the protist parasite *Theileria annulata* (Hall et al., 1990).

After removing all small contigs, and contigs belonging to canola or endophytic microbes, the final telomere-to-telomere genome assembly was at the chromosome level with an N50 of 1.37 MB, and a total size of 25.3 Mb (Figure 1, Table S1), size consistent with previous reports (Stjelja et al., 2019; Liu et al., 2023). Analysis of benchmarking universal single-copy orthologs revealed a gene completeness (BUSCO score) of 88.2%, the highest to date, with no duplications found in our final assembly (Figure 1). The genome alignment revealed a very high collinearity with the e3 strain genome, exhibiting 98.67% similarity and 1.33% of the bases in regions found to be unique between the two strains (Figure S1). The 11.88% of the complete nuclear genome, representing 3MB, was identified as repeat regions. These repeat regions are primarily composed of interspersed repeats (Gypsy, Copia, SINE, LINE) and unclassified repeats (Table S2), and are evenly distributed across all chromosomes (Figure 2A). Plotting the coding and non-coding regions on the chromosome ideogram using a 10 kb overlapping sliding window highlights long stretches of non-coding regions, indicative of putative centromeric regions (Figure 2A).

**Figure 1.**
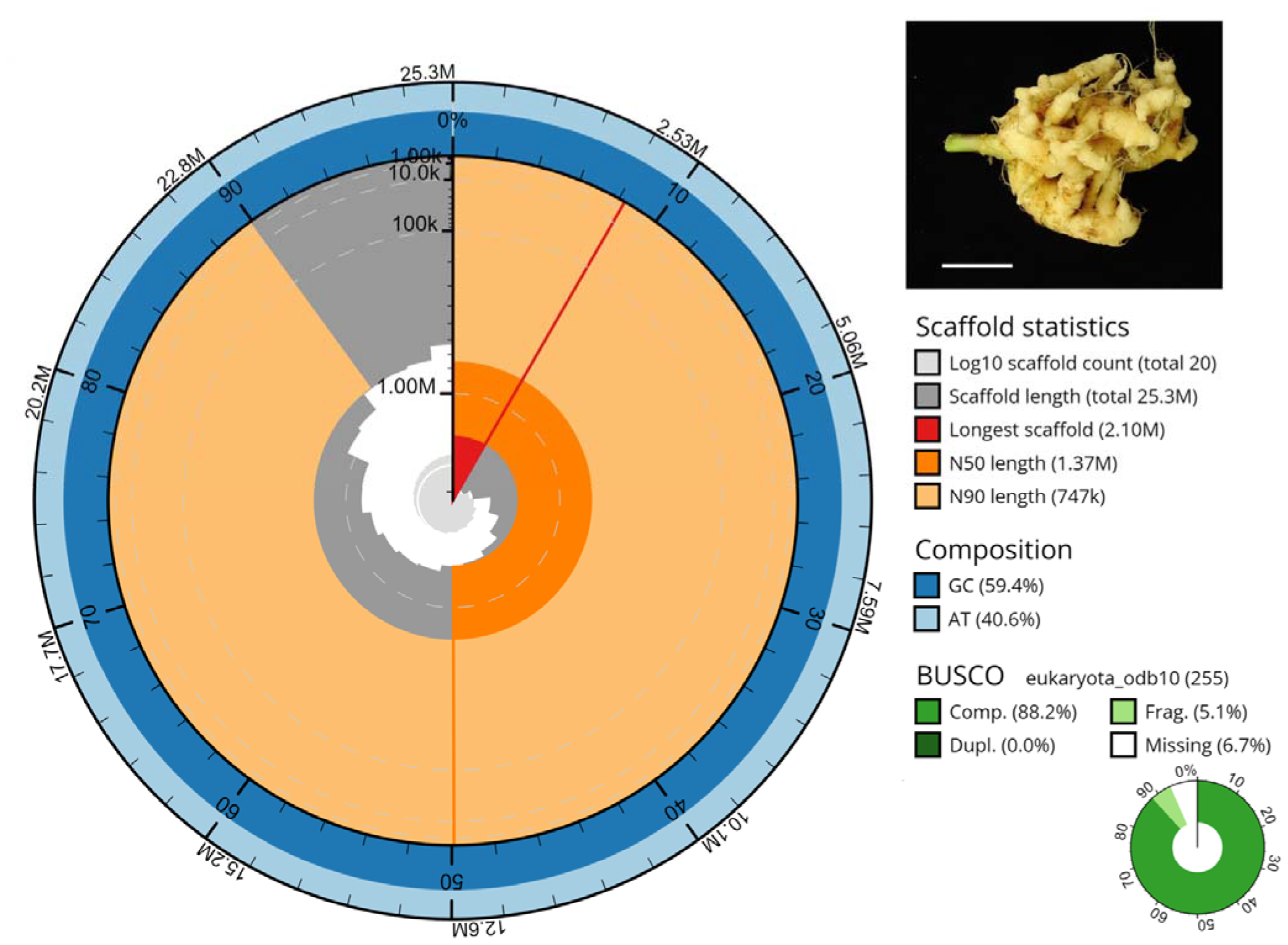
Assembly snail plot, assembly statistics, sequence composition, and BUSCO comparisons of Plasmodiophora brassicae, causal agent of clubroot disease presented on the top right panel (Bar = 1 cm).

**Figure 2.**
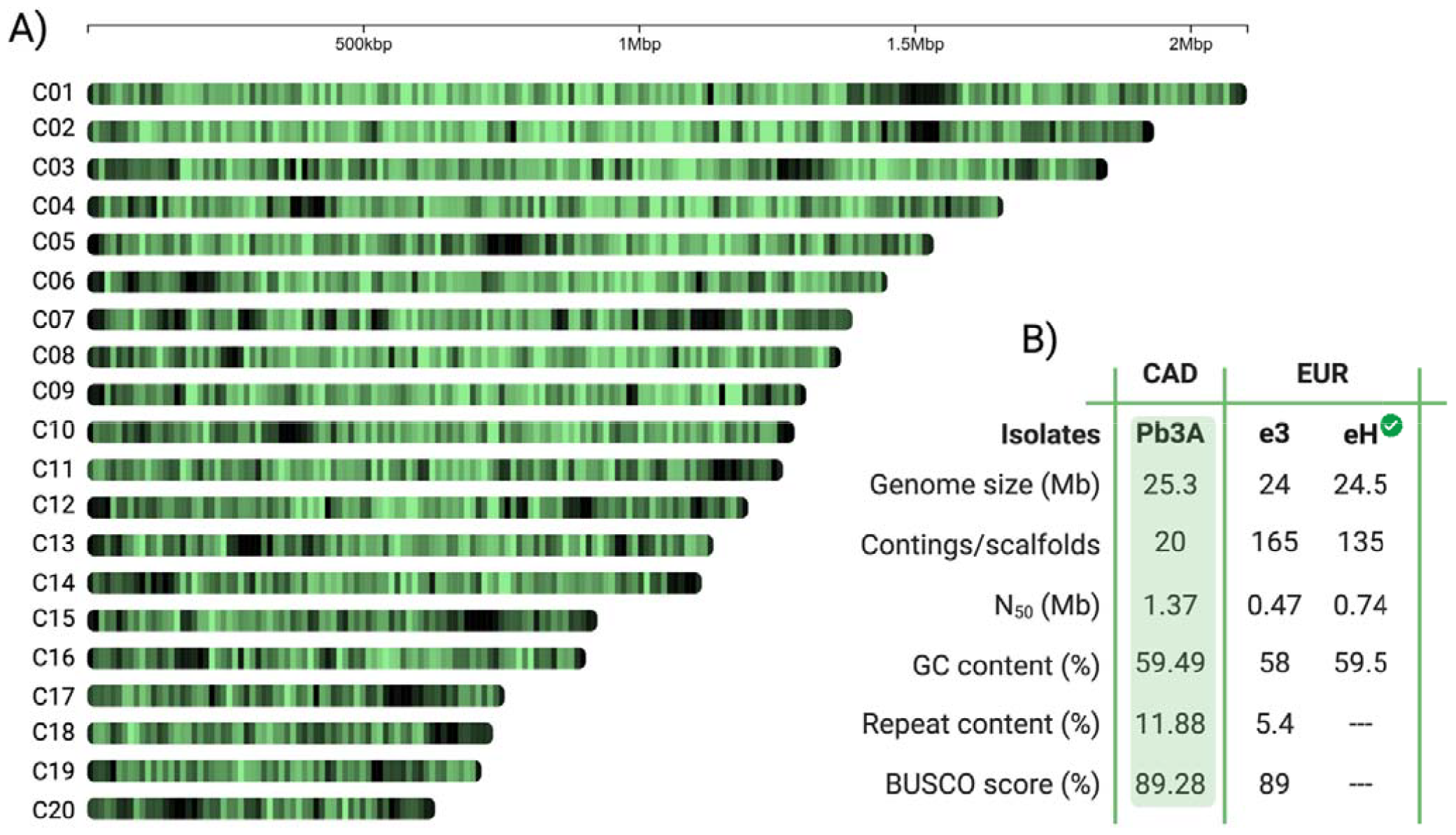
Representation and statistics of the Plasmodiophora brassicae Pb3A genome assembly. A, Ideogram of Pb3A chromosomes. Density of coding sequences is presented in green and repeat sequences in black, including telomeres at the extreme of the chromosomes, in non-overlapping sliding 10 kb windows. B, comparison of the in Pb3A genome with the first genome sequenced for the clubroot pathogen, the strain e3 and the current reference genome produced from the strain eH. BUSCO score is based on protein annotation data.

### Genome Annotation

The de-novo annotation predicted 12,590 transcripts and 10,521 protein encoding genes in the Pb3A genome assembly (Figure 2B). These values are higher than those reported for the current reference annotation in strain e3, which has 9,731 protein-encoding genes (Schwelm et al., 2015), or for the same strain later re-sequenced using long-read technology, where 9,231 protein-encoding genes were reported (Stljia et al., 2019). Overall, using our final genome assembly we annotated a higher number of protein-encoding genes than the rest of gene models currently available for the clubroot pathogen. The BUSCO score for the genome annotation, using only the longest isoforms, showed an improvement compared to previous annotations with 89.28% gene completeness and no duplication (Figure 2B). InterproScan search using Pfam, Gene3D and SUPERFAMILY database identified 70.93% of the predicted proteome with known domains (Table S3). A BLAST search (min 50% coverage, E value < 10^-05) against the entire non-redundant protein database identified 9893 sequences to have a homologue with existing P. brassicae-related sequences, while 456 sequences had no significant hit (Table S4).

## MATERIALS AND METHODS

### Sample preparation and DNA Extraction

Pb3A resting spores propagated in Westar, a susceptible *Brassica napus accession*, were extracted as previously described (Salih et al., 2023). Using 3-5 mL of spores at a concentration ranging from 108-109 spores/mL, high molecular weight (HMW) DNA was extracted from the resting spores using a modified cetyltrimethylammonium bromide (CTAB) method to avoid DNA shearing, followed by a cleaning step using the Nanobind CBB kit (PacBio, USA). The quality and concentration of extracted DNA were determined using a Qubit 4 fluorometer with dsDNA broad range (BR) Assay kit (Thermo Fisher Scientific, Canada). DNA size distribution was confirmed by running a Pippin pulse electrophoresis (Sage science, USA) and Femto Pulse gel electrophoresis (Agilent Technologies, USA) at Genome Quebec and IBIS-ULaval sequencing platforms, respectively.

### Whole Genome Sequencing

A HiFi SMRTbell® library (10 kb) was constructed and sequenced on one PacBio SMRT Cell (15h) on a PacBio Sequel II instrument at Génome Québec, Canada. In parallel, an Oxford Nanopore library was constructed (40 kb) and sequenced on a FLO-MIN106 flow cell on a GridION X5 Mk1 instrument at our institution’s sequencing platform (Plateforme d’Analyse Génomique, Institut de Biologie Intégrative et des Systèmes, Université Laval, Canada). For the Illumina sequencing, a whole genome shotgun library was prepared for paired-end 2 × 150 bp reads and sequenced on an Illumina Novaseq system at the Génome Québec sequencing facility.

### Genome Assembly

The quality of PacBio HiFi and Oxford Nanopore reads was analyzed using Qiagen CLC Genomics workbench v.23 (QIAGEN, Denmark), while the quality of Illumina paired-end reads was analyzed with FastQC v0.11.8 (Andrews 2010), and the adapters were removed using Trimmomatic v0.39 (Bolger et al., 2014). Nanopore reads shorter than 3 kb were trimmed using chopper v0.6.0 (De-Coster and Rademakers, 2023). To generate a *de novo* assembly, a hybrid assembly approach was utilized with PacBio HiFi and Oxford Nanopore reads using Hifiasm v0.19.5 with inbred mode using “-l0” flag (Cheng et al., 2021). The in-built RAVEN assembler of CLC Genomcis Workbench v23 was used to produce assembly with only Oxford Nanopore reads. The final genome assembly was polished with Hapo-G v1.2 utilizing Illumina paired-end reads with three rounds of iterations (Aury and Istace, 2021). The quality assessment of the polished assembly was done with the QUality ASsessment Tool, QUAST v2.5.0 (Gurevich et al., 2013), and the gene completeness was assessed using compleasm v0.2.5 (Huang and Li, 2023). Genomic alignment dot plot analysis was produced using D-Genies v1.5.0 (Cabanettes and Klopp, 2018).

### Genome Annotation

RepeatModeler v2.0.1 (Flynn et al., 2020) was used to construct a species-specific library of repeats and transposable elements. The custom library was used by RepeatMasker v4.1.2 (Tarailo-Graovac and Chen, 2009; Flynn et al., 2020) to soft mask the assembled genome. BRAKER3 v3.0.7 (Gabriel et al., 2023), which leverages both transcriptomic and protein evidence to automatically annotate the genome, was used to perform the annotation. For transcriptome evidence, publicly available 7-, 16-, 21-, and 26-days post inoculation (dpi) timepoints datasets were used to cover the main steps of P. *brassicae* life cycle in the susceptible host. The 7- and 21-dpi datasets (Zhou et al., 2020) were generated from samples infected with 3A pathotype, the same pathotype used in this study. However, the 16- and 26-dpi datasets (Schwelm et al., 2015) samples infected with the European isolate e3. Raw FastQ reads were directly supplied to BRAKER3 for evidence, which it internally maps to the genome using HISAT2 (Kim et al., 2019). For protein evidence, Eukaryotic OrthoDB11 (Kuznetsov et al., 2023) partition was downloaded from Griefswald University webpage, maintained by the BRAKER3 team (https://bioinf.uni-greifswald.de/bioinf/partitioned_odb11/). The current reference proteome of the P. brassicae and the proteome of three closely related Rhizaria - *Spongospora subterranean, Reticulomyxa filose and Bigelowiella natans* were downloaded from Ensembl protist and concatenated to the OrthoDB11 file. The output GTF file was parsed using a Python script (https://github.com/Gaius-Augustus/TSEBRA/blob/main/bin/get_longest_isoform.py) supplied by the BRAKER3 team to obtain only the longest isoform for each gene model. Compleasm v0.2.5 was used with the ‘eukaryota’ lineage flag to report the BUSCO score of the annotated proteome (Huang and Li, 2023). InterProScan v5.61-93.0 (Jones et al., 2014) was used to functionally annotate the proteome by searching against the Pfam, SUPERFAMILY and Gene3D databases. Omicsbox v3.1.11 was used to BLAST the protein sequences against the non-redundant (nr) protein database using the built in DIAMOND mapper (Buchfink et al., 2021).

## DATA AVAILABILITY

The Pb3A genome assembly has been deposited in the NCBI database under the accession number GCA_036867785.1 (BioProject ID PRJNA1071157). The NCBI SRA accession numbers for the raw sequencing data are as follows: PacBio HiFi data: SRX23767463, Oxford Nanopore data: SRX23767464, and Illumina data: SRX23767465. De novo annotation-related files are available on Zenodo: https://zenodo.org/records/10913168. The genome analysis workflow is also available on GitHub: https://github.com/Edelab.

## ACKNOWLEDGMENTS

We are grateful to the bioinformatics and sequencing support personal and infrastructure at IBIS, Université Laval, for the constant support throughout this project.

## AUTHOR CONTRIBUTION

Conceptualization: M.A.J., S.M. and E.P.L.; Experimental work: M.A.J. and A-S.B., Data analysis and presentation: M.A.J., S.M. and E.N.; Writing full draft, review, and editing: M.A.J., S.M., E.N. and E.P.L.; Supervision: E.P.L.

## LITERATURE CITED

Amezrou, R., Ducasse, A., Compain, J. et al. (2024). Quantitative pathogenicity and host adaptation in a fungal plant pathogen revealed by whole-genome sequencing. Nat. Commun. 15, 1933. doi: 10.1038/s41467-024-46191-1.

Andrews, S. (2010). FastQC: a quality control tool for high throughput sequence data. Available from: http://www.bioinformatics.babraham.ac.uk/projects/fastqc x(last accessed April 1st, 2024).

Aury, J.M. & Istace, B. (2021). Hapo-G, haplotype-aware polishing of genome assemblies with accurate reads. NAR Genom. Bioinform. 3(2), lqab034. doi: 10.1093/nargab/lqab034.

Badet, T., Tralamazza, S.M., Feurtey, A. & Croll, D. (2024). Recent reactivation of a pathogenicity-associated transposable element is associated with major chromosomal rearrangements in a fungal wheat pathogen. Nucleic Acids Res. 52, 1226–1242. doi: 10.1093/nar/gkad1214.

Bi, K., Chen, T., He, Z., Gao, Z., Zhao, Y., Liu, H. et al. (2019). Comparative genomics reveals the unique evolutionary status of Plasmodiophora brassicae and the essential role of GPCR signaling pathways. Phytopathol. Res. 1, 12. doi: 10.1186/s42483-019-0018-6.

Bolger, A.M., Lohse, M. & Usadel, B. (2014). Trimmomatic: a flexible trimmer for Illumina sequence data. Bioinformatics. 30(15), 2114–2120. doi: 10.1093/bioinformatics/btu170.

Buchfink, B., Reuter, K. & Drost, H.G. (2021). Sensitive protein alignments at tree-of-life scale using DIAMOND. Nat. Methods. 18, 366–368. doi: 10.1038/s41592-021-01101-x.

Cabanettes, F. & Klopp, C. (2018). D-GENIES: dot plot large genomes in an interactive, efficient and simple way. PeerJ. 6, e4958. doi: 10.7717/peerj.4958.

Cheng, H., Concepcion, G.T., Feng, X., Zhang, H. & Li, H. (2021). Haplotype-resolved de novo assembly using phased assembly graphs with hifiasm. Nat. Methods. 18, 170–175. doi: 10.1038/s41592-020-01056-5.

Ciaghi, S., Neuhauser, S. & Schwelm, A. (2018). Draft genome resource for the potato powdery scab pathogen Spongospora subterranea. Mol. Plant-Microb. Interac. 31, 1227–1229. doi: 10.1094/MPMI-06-18-0163-A.

Croll, D. (2024). Dimensions of genome dynamics in fungal pathogens: from fundamentals to applications. BMC Biol. 22:19. doi: 10.1186/s12915-023-01786-w.

Daval, S., Belcour, A., Gazengel, K., Legrand, L., Gouzy, J., Cottret, L. et al. (2019). Computational analysis of the Plasmodiophora brassicae genome: mitochondrial sequence description and metabolic pathway database design. Genomics. 111, 1629–1640. doi: 10.1016/j.ygeno.2018.11.013.

De Coster, W. & Rademakers, R. (2023). NanoPack2: population-scale evaluation of long-read sequencing data. Bioinformatics. 39, btad311. doi: 10.1093/bioinformatics/btad311.

Decroës, A., Calusinska, M., Delfosse, P., Bragard, C. & Legrève, A. (2019). First draft genome sequence of a Polymyxa genus member, Polymyxa betae, the protist vector of rhizomania. Microbiol. Resour. Announc. 8, e01509–18. doi: 10.1128/MRA.01509-18.

Flynn, J.M., Hubley, R., Goubert, C., Rosen, J., Clark, A.G., Feschotte, C. & Smit, A.F. (2020). RepeatModeler2 for automated genomic discovery of transposable element families. Proc. Natl. Acad. Sci. U S A. 117(17),9451–9457. doi: 10.1073/pnas.1921046117.

Gabriel, L., Brůna, T., Hoff, K.J., Ebel, M., Lomsadze, A., Borodovsky, M. & Stanke, M. (2023). BRAKER3: fully automated genome annotation using RNA-seq and protein evidence with GeneMark-ETP, AUGUSTUS and TSEBRA. bioRxiv. doi: 10.1101/2023.06.10.544449.

Gurevich, A., Saveliev, V., Vyahhi, N. & Tesler G. (2013). QUAST: quality assessment tool for genome assemblies. Bioinformatics. 29, 1072–1075. doi: 10.1093/bioinformatics/btt086.

Hall, R., Coggins, L., McKellar, S., Shiels, B. & Tait, A. (1990). Characterisation of an extrachromosomal DNA element from Theileria annulate. Mol. Biochem. Parasitol. 38, 253–260. doi: 10.1016/0166-6851(90)90028-K.

Huang, N. & Li, H. (2023). compleasm: a faster and more accurate reimplementation of BUSCO. Bioinformatics. btad595. doi: 10.1093/bioinformatics/btad595.

Javed, M. A., Schwelm, A., Zamani-Noor, N., Salih, R., Silvestre Vañó, M., Wu, J., González García, M., Heick, T. M., Luo, C., Prakash, P. & Pérez-López, E. (2023). The clubroot pathogen Plasmodiophora brassicae: A profile update. Mol. Plant Pathol. 24(2), 89–106. doi: 10.1111/mpp.13283.

Jones, P., Binns, D., Chang, H.-Y., Fraser, M., Li, W., McAnulla, C., McWilliam, H., Maslen, J., Mitchell, A., Nuka, G., Pesseat, S., Quinn, A. F., Sangrador-Vegas, A., Scheremetjew, M., Yong, S.-Y., Lopez, R., & Hunter, S. (2014). InterProScan 5: Genome-scale protein function classification. Bioinformatics. 30(9), 1236–1240. doi:10.1093/bioinformatics/btu03.

Kim, D., Paggi, J.M., Park, C., Bennett, C. & Salzberg, S.L. (2019). Graph-based genome alignment and genotyping with HISAT2 and HISAT-genotype. Nat. Biotechnol. 37, 907–915. doi: 10.1038/s41587-019-0201-4.

Kuznetsov, D., Tegenfeldt, F., Manni, M., Seppey, M., Berkeley, M., Kriventseva, E.V. & Zdobnov, E.M. (2023). OrthoDB v11: annotation of orthologs in the widest sampling of organismal diversity. Nucleic Acids Res. 6, 51(D1), D445–D451. doi: 10.1093/nar/gkac998.

Li, P., Lv, S., Zhang, Z., Su, T., Wang, W., Xin, X., Zhao, X., Li, X., Zhang, D., Yu, Y., Ma, T., Liu, G., Zhang, F. & Yu, S. (2023). Genome assembly of the plant pathogen Plasmodiophora brassicae reveals novel secreted proteins contributing to the infection of Brassica rapa. Hort. Plant J. doi: 10.1016/j.hpj.2023.09.001.

Liu, L., Qin, L., Zhou, Z., Hendriks, W.G.H.M., Liu, S. & Wei, Y. (2020). Refining the life cycle of Plasmodiophora brassicae. Phytopathol. 110, 1704–1712. doi: 10.1094/PHYTO-02-20-0029-R.

Malinowski, R., Truman, W. & Blicharz, S. (2019). Genius architect or clever thief – how Plasmodiophora brassicae reprograms host development to establish a pathogen-oriented physiological sink. Mol. Plant-Microbe Interact. 32, 1259–1266. doi: 10.1094/MPMI-03-19-0069-CR.

Mukhopadhyay, S., Garvetto, A., Neuhauser, S. & Pérez-López, E. (2024). Decoding the Arsenal: Protist Effectors and Their Impact on Photosynthetic Hosts. Mol. Plant-Microb. Interact. doi: 10.1094/MPMI-11-23-0196-CR.

Neuhauser, S., Kirchmair, M., Bulman, S. & Bass, D. (2014). Cross-kingdom host shifts of phytomyxid parasites. BMC Evol. Biol. 14, 1–13. doi: 10.1186/1471-2148-14-33.

Ochoa, J. C., Mukhopadhyay, S., Bieluszewski, T., Jędryczka, M., Malinowski, R. & Truman, W. (2023). Natural variation in Arabidopsis responses to Plasmodiophora brassicae reveals an essential role for Resistance to Plasmodiophora brasssicae 1 (RPB1). The Plant J. 116(5), 1421–1440. doi: 10.1111/tpj.16438.

Petracek, M.E., Lefebvre, P.A., Silflow, C.D. & Berman, J. (1990). Chlamydomonas telomere sequences are A+T-rich but contain three consecutive G-C base pairs. Proc. Natl. Acad. Sci. U S A. 87(21), 8222–6. doi: 10.1073/pnas.87.21.8222.

Rolfe, S.A., Strelkov, S.E., Links, M.G., Clarke, W.E., Robinson, S.J., Djavaheri, M. et al. (2016). The compact genome of the plant pathogen Plasmodiophora brassicae is adapted to intracellular interactions with host Brassica spp. BMC Genomics. 17, 272. doi: 10.1186/s12864-016-2597-2.

Salih, R., Brochu, A.-S., Prakash, P., Côté, J.-D., Pérez-López, E. (2024). A basic guide to the propagation and manipulation of the clubroot pathogen, Plasmodiphora brassicae. Curr. Protoc. doi: 10.1002/cpz1.1039.

Sedaghatkish, A., Gossen, B.D., Yu, F., Torkamaneh, D. & McDonald, M.R. (2019). Wholegenome DNA similarity and population structure of Plasmodiophora brassicae strains from Canada. BMC Genomics. 20, 744. doi: 10.1186/s12864-019-6118-y.

Schwelm, A., Fogelqvist, J., Knaust, A., Jülke, S., Lilja, T., Bonilla-Rosso, G. et al. (2015). The Plasmodiophora brassicae genome reveals insights in its life cycle and ancestry of chitin synthases. Sci. Rep. 5, 11153. doi: 10.1038/srep11153.

Stjelja, S., Fogelqvist, J., Tellgren-Roth, C. & Dixelius, C. (2019). The architecture of the Plasmodiophora brassicae nuclear and mitochondrial genomes. Sci. Rep. 9, 15753. doi: 10.1038/s41598-019-52274-7.

Tarailo-Graovac, M. & Chen, N. (2009). Using RepeatMasker to identify repetitive elements in genomic sequences. Curr Protoc Bioinform. 10. 1–4. doi: 10.1002/0471250953.bi0410s25.

Zhou, Q., Galindo-González, L., Manolii, V.Hwang, S.-F. & Strelkov, S.E. (2020). Comparative Transcriptome Analysis of Rutabaga (Brassica napus) Cultivars Indicates Activation of Salicylic Acid and Ethylene-Mediated Defenses in Response to Plasmodiophora brassicae. Int. J. Mol. Sci. 21, 8381. doi: 10.3390/ijms21218381.

