## Supplementary material for "Telomere-to-telomere genome assembly of the clubroot pathogen *Plasmodiophora brassicae*": Fig. S1, Table S1-S2

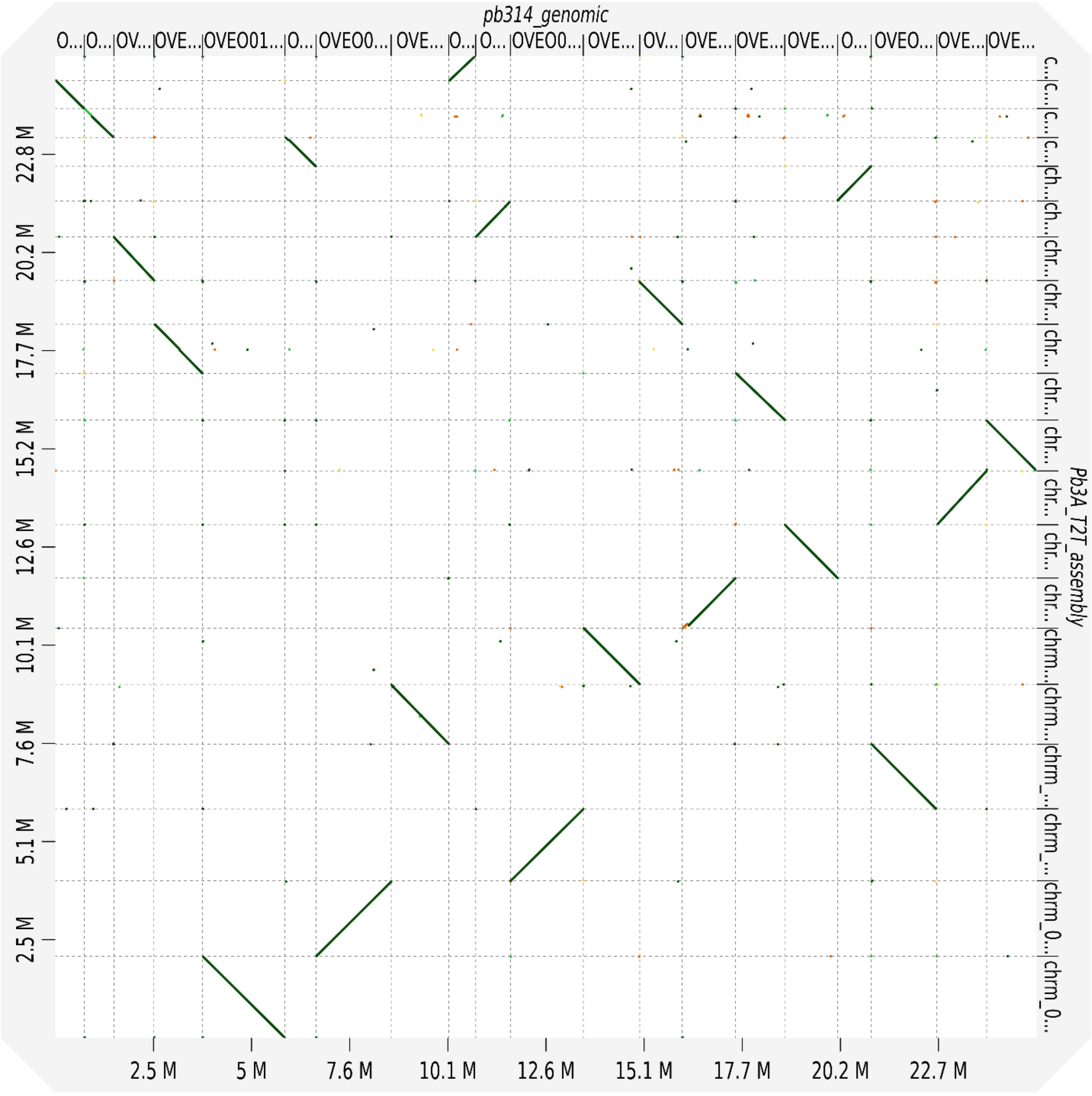


**Figure S1.** Alignment of *Plasmodiophora brassicae* Pb3A and European isolate e3 genomes using D-genies showing collinearity

**Table S1.** Chromosomes length of *Plasmodiophora brassicae* (3A) genome

| Chromosome | Size (bp) |
|---|---|
| Chrm_01 | 2102640 |
| Chrm_02 | 1943040 |
| Chrm_03 | 1855648 |
| Chrm_04 | 1659604 |
| Chrm_05 | 1544657 |
| Chrm_06 | 1445323 |
| Chrm_07 | 1389561 |
| Chrm_08 | 1371253 |
| Chrm_09 | 1307724 |
| Chrm_10 | 1291307 |
| Chrm_11 | 1260905 |
| Chrm_12 | 1206894 |
| Chrm_13 | 1127679 |
| Chrm_14 | 1123131 |
| Chrm_15 | 923546 |
| Chrm_16 | 894280 |
| Chrm_17 | 747440 |
| Chrm_18 | 741057 |
| Chrm_19 | 722821 |
| Chrm_20 | 625471 |

**Table S2.** Summary repeat element classes in *Plasmodiophora brassicae* (3A)

| Repeat Elements | Elements no. | Length occupied | Sequence % |
|---|---|---|---|
| **Retroelements** | 1871 | 1562946 bp | 6.18 % |
| SINEs | 36 | 4897 bp | 0.02 % |
| LINEs | 187 | 107712 bp | 0.43 % |
| R1/LOA/Jockey | 19 | 8643 bp | 0.03 % |
| RTE/Bov-B | 62 | 33335 bp | 0.13 % |
| LTR elements | 1648 | 1450337 bp | 5.74 % |
| Ty1/Copia | 125 | 178436 bp | 0.71 % |
| Gypsy/DIRS1 | 1005 | 1155490 bp | 4.57 % |
| **DNA transposons** | 190 | 124785 bp | 0.49 % |
| hobo-Activator | 22 | 3369 bp | 0.01 % |
| Tc1-IS630-Pogo | 92 | 61391 bp | 0.24 % |
| MULE-MuDR | 25 | 26231 bp | 0.10 % |
| **Rolling circles** | 12 | 12092 bp | 0.05 % |
| **Unclassified** | 2355 | 1044064 bp | 4.13 % |
| **Small RNA** | 9 | 29217 bp | 0.12 % |
| **Simple repeats** | 5092 | 226755 bp | 0.90 % |
| **Low complexity** | 115 | 4887 bp | 0.02 % |
| **Total** | **9,644** | **3004746 bp** | **11.88 %** |

**Note:** Most repeats fragmented by insertions or deletions have been counted as one element
Runs of >=20 X/Ns in query were excluded in % calculations
